# Pterostilbene Protects Cochlea from Ototoxicity in Streptozotocin-Induced Diabetic Rats by Inhibiting the Apoptosis

**DOI:** 10.1101/2020.01.16.908798

**Authors:** Sibel Özdaş, Bora Taştekin, Seren G. Gürgen, Talih Özdaş, Aykut Pelit, Sanem O. Erkan, Birgül Tuhanioğlu, Orhan Görgülü

## Abstract

**Objectives/Hypothesis:** Diabetes mellitus (DM) causes ototoxicity by inducing oxidative stress, microangiopathy, and apoptosis in the cochlear sensory hair cells. The natural anti-oxidant pterostilbene (PTS) (trans-3,5-dimethoxy-4-hydroxystylbene) has been reported to relieve oxidative stress and apoptosis in DM, but its role in diabetic-induced ototoxicity is unclear. This study aimed to investigate the effects of dose-dependent PTS on the cochlear cells of streptozotocin (STZ)-induced diabetic rats.

**Methods:** The study included 30 albino male Wistar rats that were randomized into five groups: non-diabetic control (Control), diabetic control (DM), and diabetic rats treated with intraperitoneal PTS at 10, 20, or 40 mg/kg/day during the four-week experimental period (DM+ PTS10, DM + PTS20, and DM + PTS40). Distortion product otoacoustic emission (DPOAE) tests were performed at the beginning and end of the study. At the end of the experimental period, apoptosis in the rat cochlea was investigated using caspase-8, cytochrome-c, and terminal deoxyribonucleotidyl transferase-mediated dUTP-biotin end labeling (TUNEL). Quantitative real-time polymerase chain reaction was used to assess the mRNA expression levels of the following genes: *CASP-3*, BCL-associated X protein (*BAX*), and *BCL-2*. Body weight, blood glucose, serum insulin, and malondialdehyde (MDA) levels in the rat groups were evaluated.

**Results:** The mean DPOAE amplitude in the DM group was significantly lower than the means of the other groups (0.9–8 kHz; P < 0.001 for all). A dose-dependent increase of the mean DPOAE amplitudes was observed with PTS treatment (P < 0.05 for all). The caspase-8 and cytochrome-c protein expressions and the number of TUNEL-positive cells in the hair cells of the Corti organs of the DM rat group were significantly higher than those of the PTS treatment and control groups (DM > DM + PTS10 > DM + PTS20 > DM + PTS40 > Control; P < 0.05 for all). PTS treatment also reduced cell apoptosis in a dose-dependent manner by increasing the mRNA expression of the anti-apoptosis *BCL2* gene and by decreasing the mRNA expressions of both the pro-apoptosis *BAX* gene and its effector *CASP-3* in a dose-dependent manner (P < 0.05 compared to DM for all). PTS treatment significantly improved the metabolic parameters of the diabetic rats, such as body weight, blood glucose, serum insulin, and MDA levels, consistent with our other findings (P < 0.05 compared to DM for all).

**Conclusion:** PTS decreased the cochlear damage caused by diabetes, as confirmed by DPOAE, biochemical, histopathological, immunohistochemical, and molecular findings. This study reports the first in vivo findings to suggest that PTS may be a protective therapeutic agent against diabetes-induced ototoxicity.

## Introduction

Diabetes mellitus (DM) is a common metabolic disorder that affects the metabolism of proteins, fats, and carbohydrates. DM is caused by the pancreas’s failure to secrete insulin and/or the deterioration of the tissue’s response to insulin.^1^ Globally, diabetes is currently projected to affect 425 million people, and that number that is expected to reach 629 million by 2045.^2^ Diabetes causes complications in many organs, such as the kidneys, liver, nervous system, reproductive system, and eyes.^3–5^ Type 1 DM is an autoimmune disease that is characterized by the destruction of immune-mediated beta-cells that leads to an absolute insulin deficiency.^6^ Type 2 DM is characterized by hyperglycemia, insulin resistance, and relative insulin deficiency.^7^ The insensitivity to insulin that occurs due to insulin resistance, reduced insulin production, and pancreatic beta-cell failure is specific to Type 2 DM.^8^

Sudden or gradual hearing loss has been observed in DM patients.^4,5,9^ DM has been reported to cause various histological changes and hearing loss in the cochlea, VIII nerve, and the temporal bone.^10,11^ The distortion product otoacoustic emission (DPOAE) is a low-level sound that can be measured in the external ear canal as a reflection of active processes in the cochlea.^12^ In diabetic patients, DPOAE amplitudes have been reported to decrease functionally, especially at high frequencies.^13^

In DM, metabolic stress and oxygen reduction have been shown to increase free oxygen radical production and to decrease the protective capacity of the antioxidant defense system.^14^ DM contributes to the development of complications such as blindness, heart-vessel disease, and kidney disease, leading to a series of cascade reactions that cause the excessive production of free radicals.^15^

Pterostilbene (PTS) (trans-3,5-dimethoxy-4-hydroxystylbene) is a natural antioxidant derived from the *Pterocarpus marsupium* (leguminasae) tree, which is also known as Indian kino or bijasar.^16^ *P. marsupium* has been used traditionally in public medicine to treat diabetes and has been shown to control blood sugar levels in diabetic experimental animals.^17^ PTS is a natural methoxylated derivative of resveratrol.^18^ DM animal model studies have shown that PTS treatment significantly increased the production of endogenous antioxidants and decreased blood sugar levels.^17,19^ This study aimed to investigate the effects of dose-dependent PTS on the cochlear cells of streptozotocin (STZ)-induced diabetic rats.

## Materials and Methods

### Animals and Care

The study included 30 albino male Wistar rats that had an average weight of 250–300g. All the animals were housed in separate cages, in conditions of 50–60% humidity and 22 ± 2 C° heat, and fed standard pellet feed (in the form of seasonal fresh vegetables and fruits) and tap water. The lighting in the room was rotated between 12 hours bright and 12 hours dark. All the rats underwent otoscopic examinations, and only the animals who passed the Distortion Product Otoacoustic Emission (DPOAE) test were included in the study. Rats with ear disease, a tympanic membrane anomaly, or who did not pass the DPOAE test were excluded from the study.

### Ethics Statement

The study was handled according to the Public Health Service (PHS) Policy on Humane Care and Use of Laboratory Animals, the Animal Welfare Act, and the NIH’s guidelines. The study protocol was approved by the Ethics Committee of Celal Bayar University, Manisa, Turkey (ID: 700/2019).

### Chemicals

STZ, ethylenediamine tetra-acetic acid (EDTA), dimethyl sulfoxide (DMSO), sodium citrate, and phosphate buffered saline (PBS) were purchased from Sigma-Aldrich (St Louis, MO, USA). PTS was supplied as a free sample from Sabinsa Corporation, USA, and its 99% purity was determined by high performance liquid chromatography (HPLC).^20^

### STZ-Induction of DM

The STZ rats were starved for 16 hours, and their water intake was restricted. Fresh STZ was prepared in a cold 20 mM solution of sodium citrate (pH 4.5). Within 15 minutes of the preparation, 45 mg/kg (a single dose) of STZ was administered intravenously (IV) through the tail vein of each rat.^21^ After 72 hours, tail vein blood samples were obtained from the rats. The glucose levels of the samples were ≥ 300 mg/dL (16.7 mmol/l), indicating that the rats were considered diabetic.^22^ However, any rats with excessive weight loss, weak body resistance, or blood glucose levels above 500 mg/dL were excluded from the study; any rats that were ill or deceased were also excluded from the study.^23^

### Experimental Design

This study used 30 rats (6 normal control rats and 24 diabetic rats). After STZ-induced DM was established, the rats were divided into five groups, with each group containing 6 rats. PTS was administered intraperitoneally (IP) each day to the diabetic rats in a buffer solution that contained 10% DMSO. The study design is provided below.

Control (*n* = 6); non-diabetic control rats treated with 0.1 M sodium citrate buffer, IV) DM (*n* = 6); diabetic control rats treated with STZ (45 mg/kg/day bodyweight, IV) DM+PTS10 (*n* = 6); diabetic rats treated with PTS (10 mg/kg/day bodyweight, per day, IP) DM+PTS20 (*n* = 6); diabetic rats treated with PTS (20 mg/kg/day bodyweight, per day, IP) DM+PTS40 (*n* = 6); diabetic rats treated with PTS,40 mg/kg/day bodyweight, per day, IP)

### DPOAE Test

DPOAE are low-level acoustic signals that can be measured from the ear canal in the presence of an external acoustic stimulus.^24^ All the rats were twice tested for DPOAE, both at the beginning of the study and at the end of the 4-week experimental period. Sodium pentobarbital (75 mg/kg; Akorn, Lake Forest, IL, United States) was administered IP for sedation before each measurement. The DPOAE were measured using a Neuro-Audio/OAE device (version 2010, Neurosoft, Ivanovo, Russia) that was set at 500–8,000 Hz and recorded in DP gram. The DPOAE stimulus intensity was set to 55 for the L1 and L2 levels; the ratio of the f1 frequency (65 dB SPL) and the f2 frequency (55 dB SPL) (f2/f1) was set to 1.22. The signal-to-noise ratio values were recorded at different frequencies (0.988, 2.222, 2.963, 5.714, and 8.000 kHz).

### Anesthesia and Tissue Preparation

After the DPOAE test was repeated at the end of the 4-week experimental period, all the animals were anesthetized with a combination of ketamine hydrochloride (15 mg/kg; Pfizer, Walton Oaks, United Kingdom) and xylazine hydrochloride (5 mg/kg; Bioveta, Komenskeho, Czech Republic), decapitated, and sacrificed. The cochleas were removed from the temporal bones and dissected in cold PBS. For the histological examination, the cochleas were preserved with neutral formalin.

### Histopathological Examination

The cochlear tissues were fixed in neutral formalin for 24 hours. The tissues were then placed in a 0.1 mol/L EDTA solution to decalcify the osseous portions. This procedure was followed by an overnight washing under flowing water. The tissues were dehydrated with an alcohol series, and transparency was achieved using xylene. Serial sections (5 µm in thickness) were mounted to polylysine-coated slides. Images were obtained from the basal turn of each cochlea.

### Immunohistochemistry

For immunohistochemical staining, the remaining serial sections were assigned, incubated at 60°C overnight, and the slides were deparaffinized by xylene and dehydrated through an alcohol series. The sections were boiled for 15 min in citrate buffer (10 mM, pH 6.0) using a microwave oven to retrieve antigen. Furthermore, for preventing endogenous peroxidase activity, they were placed in hydrogen peroxidase for 15 min The sections were incubated in a blocking serum (Ultra V Block, TP-060-HL; NeoMarker, Fremont, CA) for 10 min and then with primary antibodies, including Caspase-8 (1:100, LabVision, USA) and Cytocrome c (1:100, LabVision, USA) in a moist environment at room temperature for 60 min. The antigen-antibody complex was fixed with biotinylated secondary antibodies and streptavidin-peroxidase for 20 min. Labeling was performed using DAB, and background staining was achieved using Mayer’s hematoxylin and covered by mounting medium. The images were captured using a camera attached to an Olympus microscope (CX31, Germany). Except for omission of the incubation period with the primary antibody, control samples were similarly processed. Two pathologists, blinded to the study, independently evaluated the immunolabeling scores. Scores of the staining intensity of the slides was semi-quantitatively assigned. The staining intensity was decided weak, moderate, or strong and valued 1, 2, or 3, respectively. The score was obtained from the following: SCORE=ΣPi (i+1), where i is the intensity of staining, and Pi is the percentage of stained cells for each intensity, varying from 100% to 0%.

### Terminal Deoxribonucleotityl transferase-mediated dUTP-biotin end Labeling (TUNEL)

A specific kit was used for detecting DNA fragmentation and apoptotic cell death *in situ* (Apoptag, S7101, Chemicon, CA, USA). Sections were stored at 60°C heat overnight in an oven in order to facilitate deparaffinization. To complete deparaffinization, the sections were reacted with xylol for 2 rounds of 15 min each. The sections were then placed into an alcohol series of 100%, 96%, and 80% by 10-min intervals. After 2 washings in distilled water for 5 min, the sections were incubated with 20 μg/mL proteinase K. Next, the endogen peroxidase activity was blocked in the sections, and they were allowed to react with 3% hydrogen peroxide (TA-015- HP; Lab Vision, Fremont, CA) after washing with PBS. Subsequently, the sections were incubated in a balanced buffer for 15 min and then incubated with a TdT enzyme (77 μL reaction buffer+33 μL TdT enzyme, 1-μL TdT enzyme) at 37°C for 60 min. Further, the sections were placed in pre-warmed stopping/washing buffer at room temperature for 10 min before incubating them with anti-digoxigenin for 45 min. PBS washing was performed after every step. After the washing, DAB staining was used for detecting TUNEL-positive cells. For background staining, methyl green was applied for 5 min. The stained slides were dehydrated using an alcohol series and placed in xylol for 20 min. The slides were then covered by entellan and thin glass. Finally, the slides were observed using a photolight microscope equipped with a computer. Two pathologists, who were blinded to the experiment, independently evaluated the TUNEL scores. The average number was determined by counting the TUNEL-positive apoptotic cells in randomly selected fields of each case. A total of 100 TUNEL-positive or - negative cells were counted in each case, and the TUNEL-positive cells were provided as a percentage. The cells in necrotic regions and those having a poor morphology and borders between sections were excluded.

### Quantitative Real-Time Polymerase Chain Reaction (qRT‐PCR)

The cochlear tissue was dissected in cold-PBS and homogenized with the Qiazol lysis reagent (Qiagen, Hilden, Germany). A high pure RNA isolation kit (Roche Diagnostic, GmbH, Germany) was used to isolate the total RNA in the homogenate according to the manufacturer’s protocol. The RNA concentration was evaluated with a spectrophotometer (Shimadzu UV-mini 1240, GmbH, Germany). The complementary DNA (cDNA) was synthesized using a transcriptor high fidelity cDNA synthesis kit (Roche Applied Science, Penzberg, Germany). Each prepared sample used 20 µL of the SYBR green qPCR reaction kit (Roche Applied Science), which contained 2 µL of cDNA for qRT‐PCR using the following primer pairs: *CASP-3*, forward 5’-GGAGCAGCTTTGTGTGTGTG and reverse 5’- CTTTCCAGTCAGACTCCGGC-3’; *BAX*, forward 5’-GTTTCATCCAGGATCGAGCAG-3’ and reverse 5’-CATCTTCTTCCAGATGGT GA-3’; *BCL-2*, forward 5’- CCTGTGGATGACTGAGTACC-3’ and reverse 5’-GAGACAGCCAGGAGAAATCA-3’; GAPDH, forward 5’-CTTCCGTGTTCCTACCCCCAATGT-3’ and reverse 5’- GCCTGCTTCACCACCTTCTTGATG-3’. The *GAPDH* sequences were used for normalization, as it represents a housekeeping gene. The rotor-gene Q 5plex HRM platform (Qiagen, Hilden, Germany) was used for the qRT-PCR, and the data were analyzed using the comparative Ct method (^∆∆^C_T_).

### Body Weight and Biochemical Investigations

During the experiment, the animals’ blood glucose levels and body weights were measured once a week.^25^ The blood samples were taken from the rats’ tail veins. The blood glucose levels were measured using a glucometer (Accu-Chek Performa Nano, Roche Diagnostics, Mannheim, Germany). The rats’ insulin levels were measured using an enzyme-linked immunosorbent assay (ELISA) kit (Hangzhou Eastbiopharm Co., Ltd., Hangzhou, China) and a spectrophotometer (Biotek Epoch™ Microplate Spectrophotometer, United States). Malondialdehyde (MDA) levels were measured at an absorbance of 532 nm (Shimadzu UV-mini 1240, GmbH, Germany).^26^

### Statistical Analysis

The Statistical Package for the Social Sciences (SPSS version 20.0, IBM, Inc., Chicago, IL, United States) software program was used to analyze the data. The data from the DPOAE tests, the TUNEL assays, the immunohistochemical staining, and the biochemical tests were compared between the groups with the one-way analysis of variance (ANOVA) test, followed by Tukey’s post hoc test. The Mann–Whitney U test was used to compare the means of the groups for the qRT-PCR data. All the data are presented as the mean ± the standard error of the mean. P values < 0.05 were considered statistically significant.

## Results

### DPOAE Measurements

The DPOAE amplitudes were recorded, and the averages were compared against the 0.9–8 kHz frequencies of each rat group (Table 1). The means of the DPOAE amplitudes in the DM group were significantly lower at all frequencies compared to both the Control group and the other treatment groups (P < 0.001 for all). In all the DM + PTS treatment groups, a significant increase was observed in the DPOAE amplitude averages at every frequency compared to the untreated diabetic rats (P < 0.05 for all). The highest amplitude increase was observed in the diabetic rats that received 40 mg/kg/day of PTS (P = 0.05, P = 0.001, P = 0.003, P < 0.001, P < 0.001, respectively). In addition, a significant increase was observed in the mean DPOAE amplitudes compared to the other diabetic treatment groups that received PTS treatments of 20 and 40 mg/kg/day compared to the mean of the DPOAE amplitudes at every frequency (P < 0.05 for all). However, the mean values of the DPOAE amplitudes of the healthy Control group animals were similar to those of the diabetic rats that received the PTS treatment of 40 mg/kg/day at each frequency (P = 0.998, P = 0.995, P = 0.998, P = 0.914, P = 0.934, respectively).

**Table 1.** The DPOAE amplitude (dB SPL) measurement results of the rat groups

| **Groups** | **0.988 kHz** | **2.222 kHz** | **2.963 kHz** | **5.714 kHz** | **8.000 kHz** |
| --- | --- | --- | --- | --- | --- |
| **Control (n = 6)** | 5.80 ± 4.1 | 5.83 ± 3.5 | 5.13 ± 2.4 | 10.28 ± 5.4 | 12.63 ± 9.7 |
| **DM (n = 6)** | 4.83 ± 5.6 | 4.28 ± 3.3 | 4.20 ± 3.6 | 7.66 ± 5.0 | 9.63 ± 5.5 |
| **DM+PTS10 (n = 6)** | 6.68 ± 5.8 | 5.63 ± 5.7 | 6.37 ± 4.5 | 10.28 ± 4.7 | 12.73 ± 5.7 |
| **DM+PTS20 (n = 6)** | 6.35 ± 5.9 | 5.46 ± 0.1 | 5.41 ± 6.4 | 10.19 ± 5.2 | 12.57 ± 4.4 |
| **DM+PTS40 (n = 6)** | 5.70 ± 4.9 | 5.91 ± 2.6 | 5.20 ± 2.2 | 10.52 ± 4.2 | 12.93 ± 6.8 |
| ***P value*** | *0.000* | *0.000* | *0.000* | *0.000* | *0.000* |

DPOAE: The Distortion Product Otoacoustic Emission; DM: Diabetes Mellitus; PTS: Pterostilbene

### Immunohistochemical and Histopathological Examinations

The cells in all the study groups were stained and counted using the caspase-8, cytochrome-c, and TUNEL assays (Figure 1).

**Fig1.**
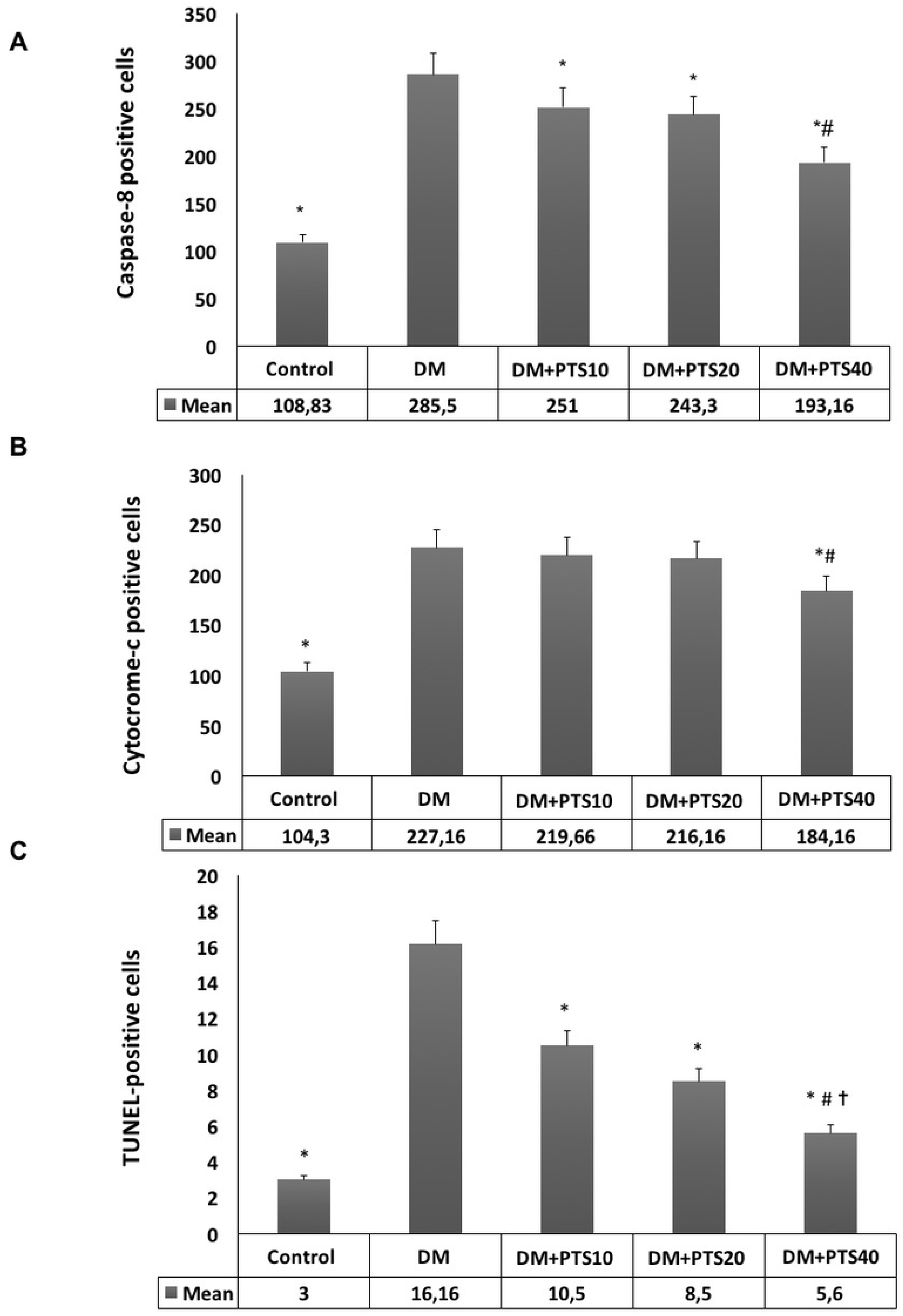
The cells counted staining using Caspase-8, Cytocrome-c, and TUNEL method in the rat groups after STZ-application to end of the experimental period. (A). For Caspase-8; ∗significantly different from DM (P< 0.001 for all), # different from DM+PTS10, DM+PTS20 (P< 0.001 for all). (B) For Cytocrome-c; ∗ Significantly different from DM (P< 0.001 for all), # different from DM+PTS10, DM+PTS20 (P< 0.001 for all). (C) For Tunnel-method; ∗ Significantly different from DM (P< 0.05 for all), # different from DM+PTS10 (P= 0.033), and✝similar to Control.

While poor expression was observed in the Control group’s Corti organ in the caspase-8 immune staining (Figure 2A), significant staining was noted in the DM group (Figure B). A moderate-to-severe caspase-8 immunoreaction was observed in the Corti organ of the diabetic rats that had been treated with PTS at 10 mg/kg/day (Figure 2C). In the group of diabetic rats that had been treated with PTS at 20 mg/kg/day, a moderate caspase-8 immunoreaction was observed (Figure 2D). Poor expression was detected in the diabetic rats that were treated with PTS at 40 mg/kg/day (Figure 2E).

**Fig2.**
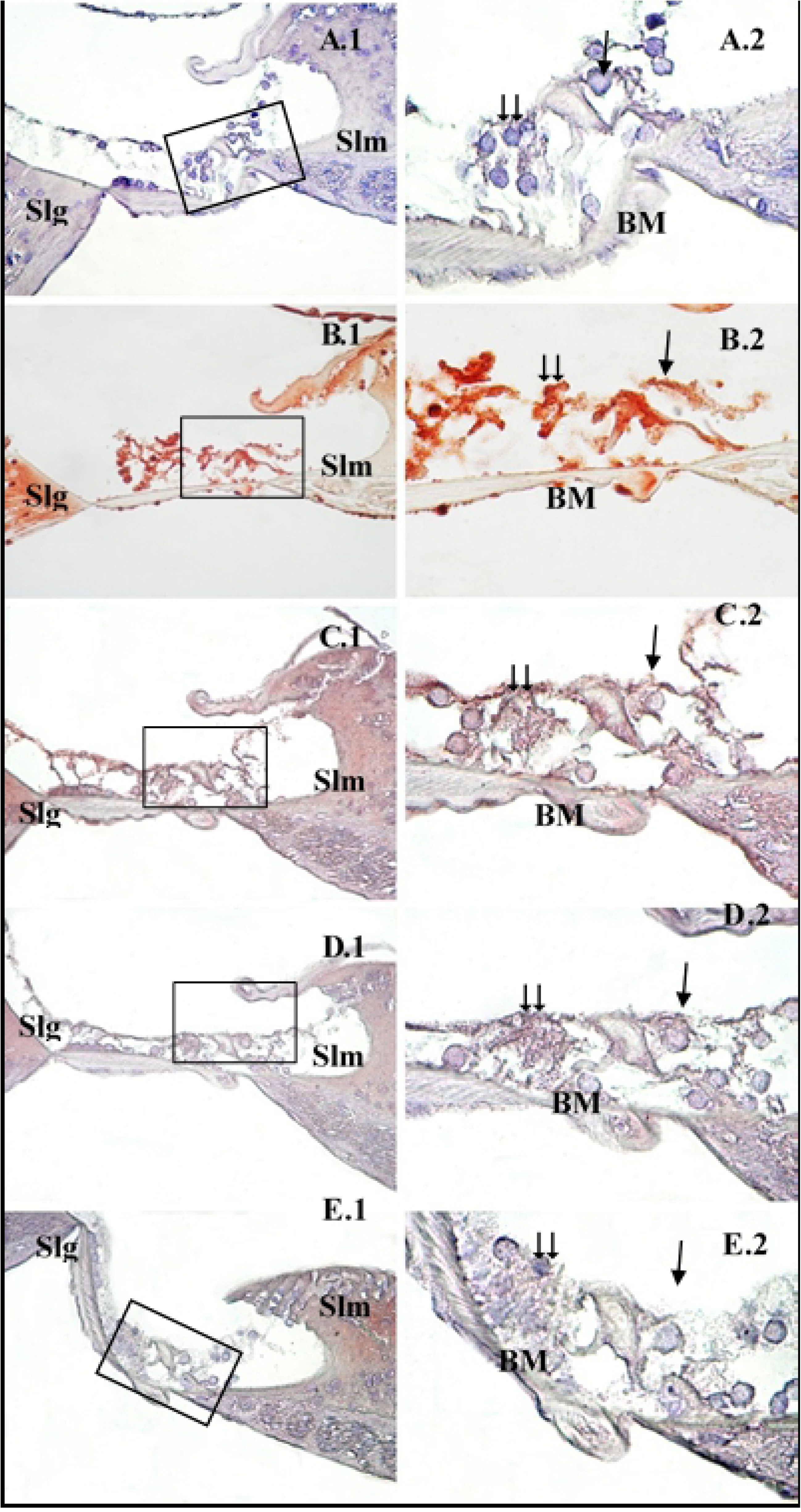
Caspase-8 immunohistochemistry staining in the cochlear Corti organ of the rat groups.
**→**: İnner hair cells, **⇉**: Outer hair cells, **Slm**: Spiral limbus, **Slg**: Spiral ligament, **BM**: Basilar membrane, Control (A), DM (B), DM+PTS10 (C), DM+PTS20 (D), DM+PTS40 (E). Corti organ X40 (1), Corti organ X100 (2). Methyl-green ground staining. X100 (2). Methyl-green ground staining.

The cytochrome-c immunodeficiency dye had a weak reaction in the Corti organs of the healthy Control group (Figure 3A). In the DM group, an increased dye reaction was observed in the Corti organ, especially in the outer hair cells, which ranged from medium to high reaction levels (Figure 3B). In the diabetic rats that were treated with PTS at 10 and 20 mg/kg/day, cytochrome-c expression ranged from medium to weak in the Corti organ cells (Figure 3C– 3D). Weak staining was observed in the diabetic rats treated with PTS at 40 mg/kg/day (Figure 3E).

**Fig3.**
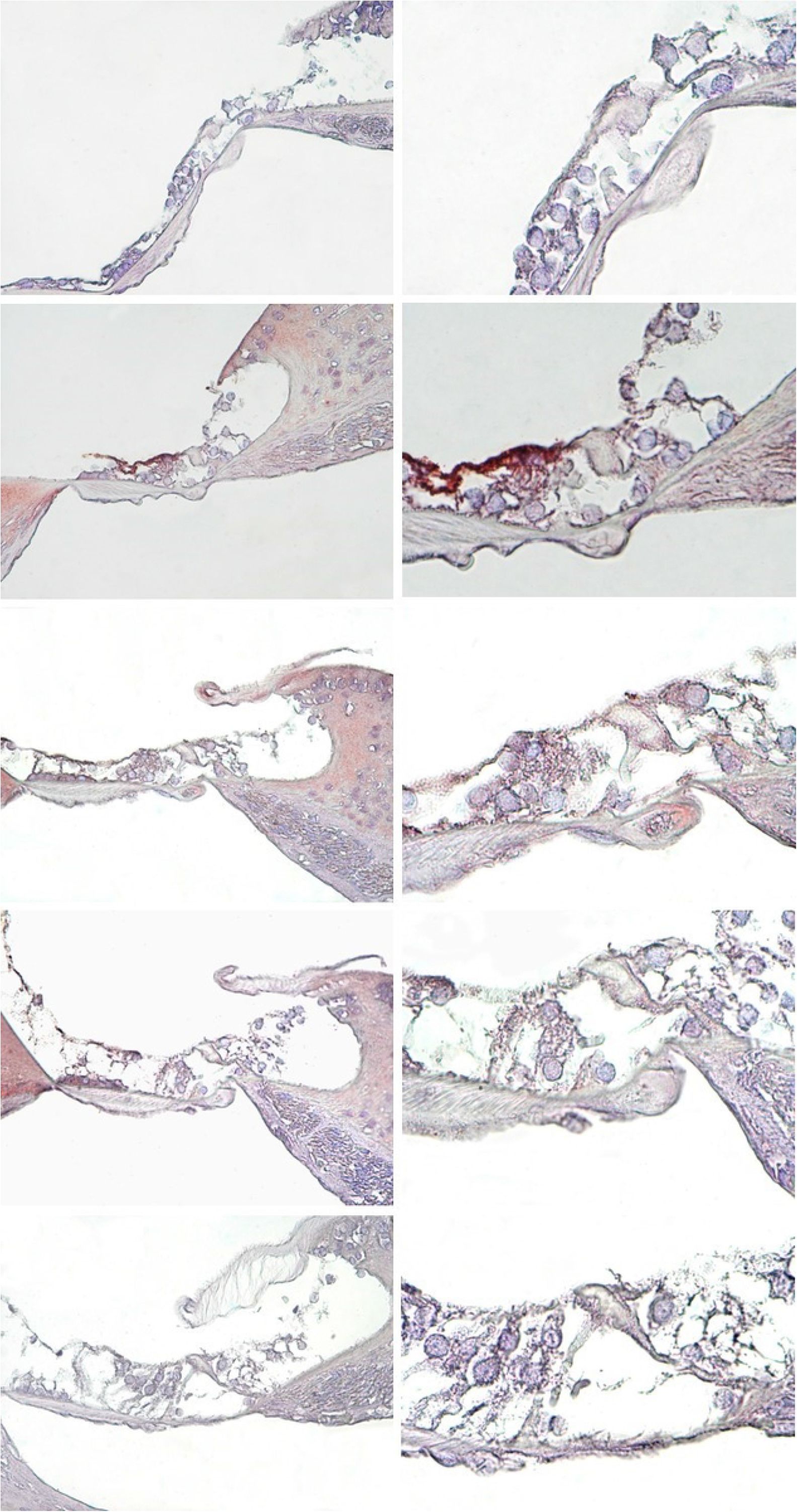
Cytocrome-c immunohistochemistry staining in the cochlear Corti organ of the rat groups. **→**: İnner hair cells, **⇉**: Outer hair cells, **Slm**: Spiral limbus, **Slg**: Spiral ligament, **BM**: Basilar membrane, Control (A), DM (B), DM+PTS10 (C), DM+PTS20 (D), DM+PTS40 (E). Corti organ X40 (1), Corti organ X100 (2). Methyl-green ground staining.

In the tissues from the healthy Control group that were stained using the TUNEL assay, there were few or no TUNEL-positive cells in the Corti organ (Figure 4A). In the DM group, strong TUNEL reactions were seen in the outer hair cells of the Corti organ, as well as the other supporting cells of the Corti organ (Figure 4B). In the diabetic rats treated with PTS at 10 mg/kg/day, fewer outer hair cells had TUNEL reactions, while TUNEL-positive reactions were observed in the support cells (Figure 4C). In the diabetic rats that were treated with PTS at 20 mg/kg/day, the outer hair cells were TUNEL-negative, and the support cells were partially TUNEL-positive (Figure 4D). In the diabetic rats that were treated with PTS at 40 mg/kg/day, the TUNEL reactions were minimal in both the outer hair cells and the support cells (Figure 4E).

**Fig4.**
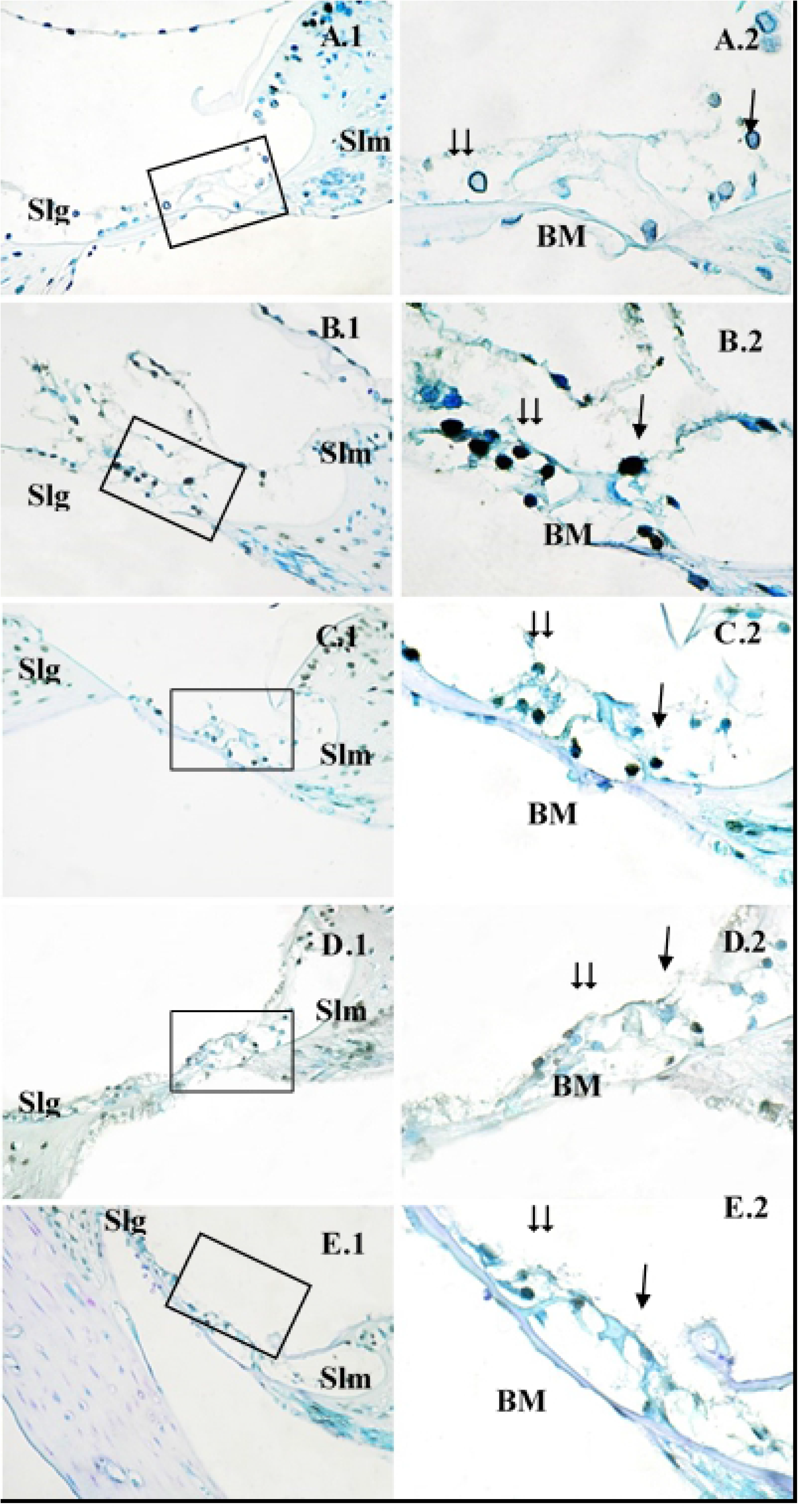
TUNEL staining in the cochlear Corti organ of the rat groups. **→**: Inner hair cells, **⇉**: Outer hair cells, **Slm**: Spiral limbus, **Slg**: Spiral ligament, **BM**: Basilar membrane, Control (A), DM (B), DM+PTS10 (C), DM+PTS20 (D), DM+PTS40 (E). Corti organ X40 (1), Corti organ

### The mRNA Expressions of the CASP3, BAX and BCL2 genes by qRT‐PCR

The mRNA levels of the *BAX, CASP-3* and *BCL-2* genes were assessed with qRT-PCR to investigate whether PTS affected apoptosis (Figure 5). In the untreated diabetic rats, compared to the rats in the healthy Control group and all the PTS treatment groups, the mRNA levels of the *BAX* and *CASP-3* genes were significantly higher (P < 0.001 for all). When the diabetic rats that received PTS were compared to the untreated diabetic rats, the mRNA levels of the *CASP-3* and *BAX* genes of the rats that received 20 and 40 mg/kg/day of PTS were significantly lower, and the *BCL-2* levels were higher (P < 0.05 for all) (Figure 5).

**Fig5.**
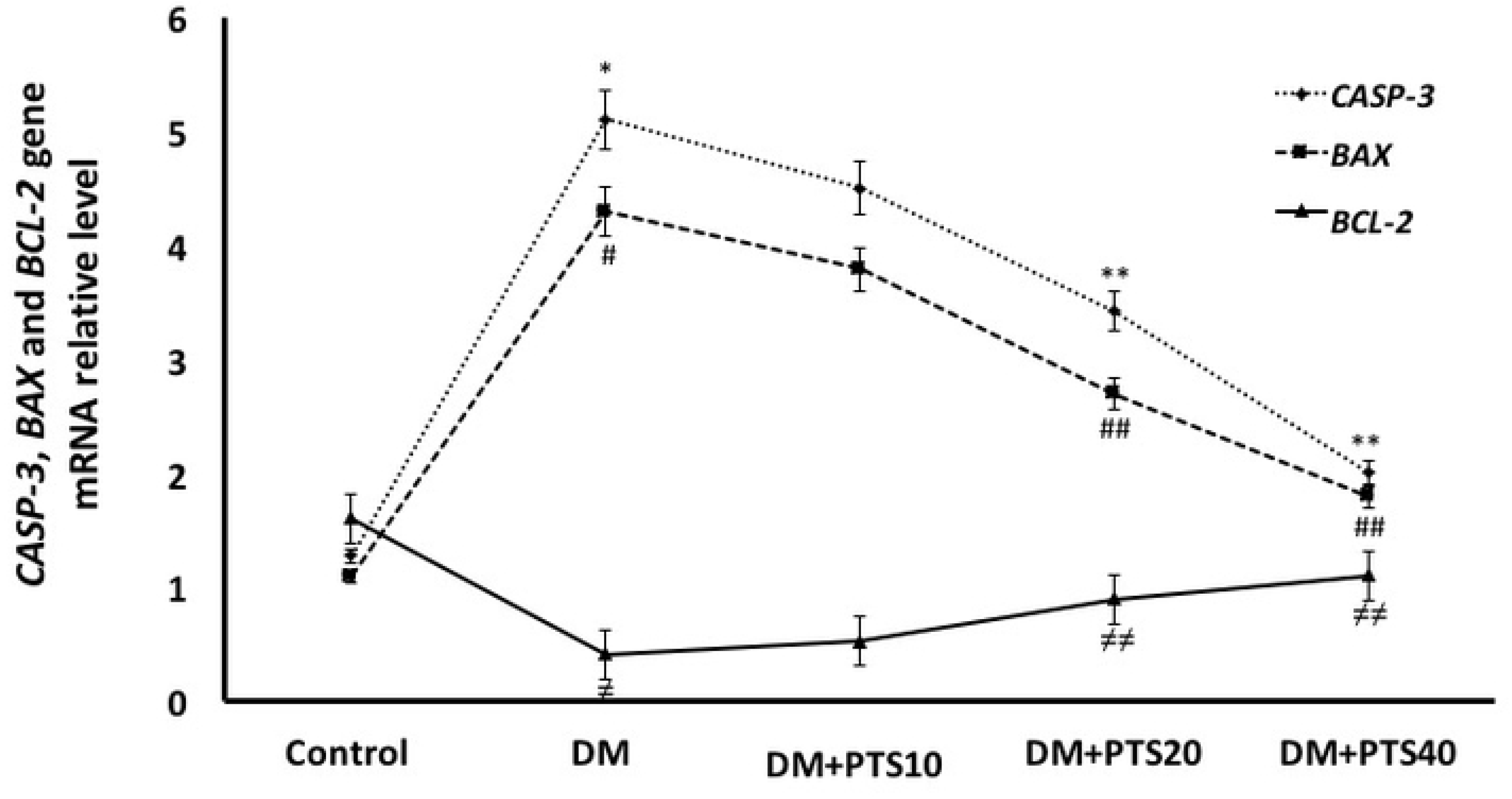
Effect of PTS on *CASP-3, BAX* and *BCL-2* gene expressions in the rat groups after STZ-application to end of the experimental period. ∗, #, ≠ significantly different from Control and treatment rat groups (P< 0.001 for all); ‪*, ##, ≠≠ significantly different from DM+PTS10 (P< 0.05 for all).

### Body Weight and Biochemical Measurements

The rats were evaluated for weight loss during the 4-week experiment. Significant weight loss was observed in the untreated diabetic rats compared to the healthy Control group and the diabetic groups that received PTS (P < 0.001). While weight gain was observed in the PTS treatment groups compared to the untreated diabetic rats, no significant differences were observed between the rats that were treated with PTS at 10 and 20 mg/kg/day (P = 0.978, P = 0.514). However, a significant weight gain was detected in the group that received 40 mg/kg/day of PTS compared to the groups that received the lower doses, and similarities were observed between the group that received the highest PTS level and the Control group (P < 0.001 for the other doses, P = 0.822 for the Control group) (Figure 6).

The rats’ blood glucose values were measured after the STZ application. 3. Daily measurements were taken during the first week and were continued until the end of the fourth week. The blood glucose values were then compared between the rat groups. The blood glucose values of the rat groups were reported with ± standard error of values on average, and a statistically significant difference was observed between all groups (P < 0.001 for all). The blood glucose levels were significantly higher in the diabetic rats that received PTS treatment compared to the healthy Control group (P < 0.001). A significant decrease in blood glucose levels was observed in the rats that were treated with 20 and 40 mg/kg/day of PTS compared to the untreated diabetic rats (P = 0.003, P < 0.001, respectively). In addition, the diabetic rats that received 20 and 40 mg/kg/day of PTS had similarly reduced blood glucose levels, and their blood glucose levels were lower than the diabetic rats that received 10 mg/kg/day of PTS (P = 0.038 and P < 0.001 for others) (Figure 7).

**Fig6.**
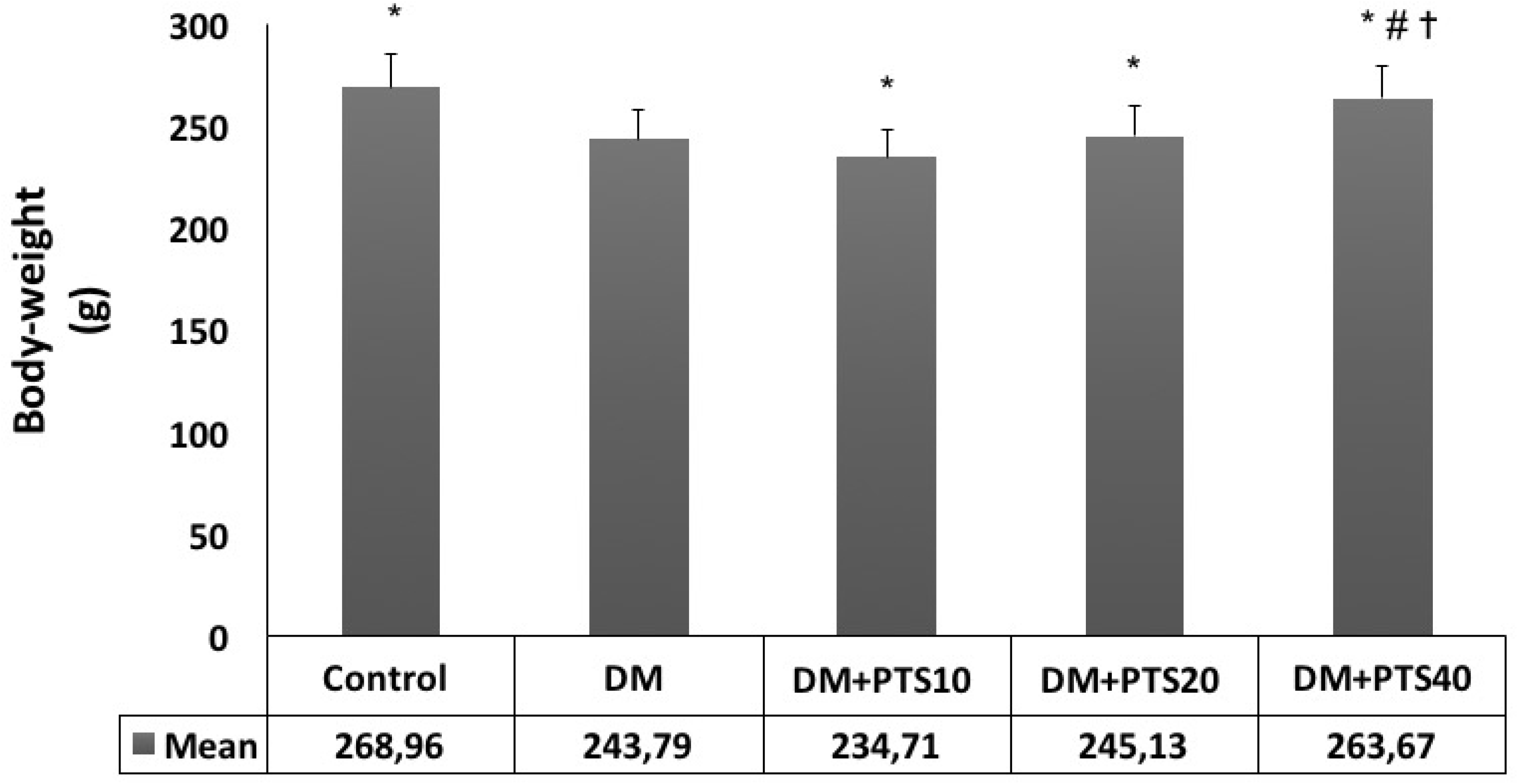
Body-weight (g) change of the rat groups after STZ-application to end of the experimental period. ∗Significantly different from DM (P< 0.001 for all), # different from all treatment rat groups (P< 0.001 for all), and ✝similar to Control (P= 0.822).

**Fig7.**
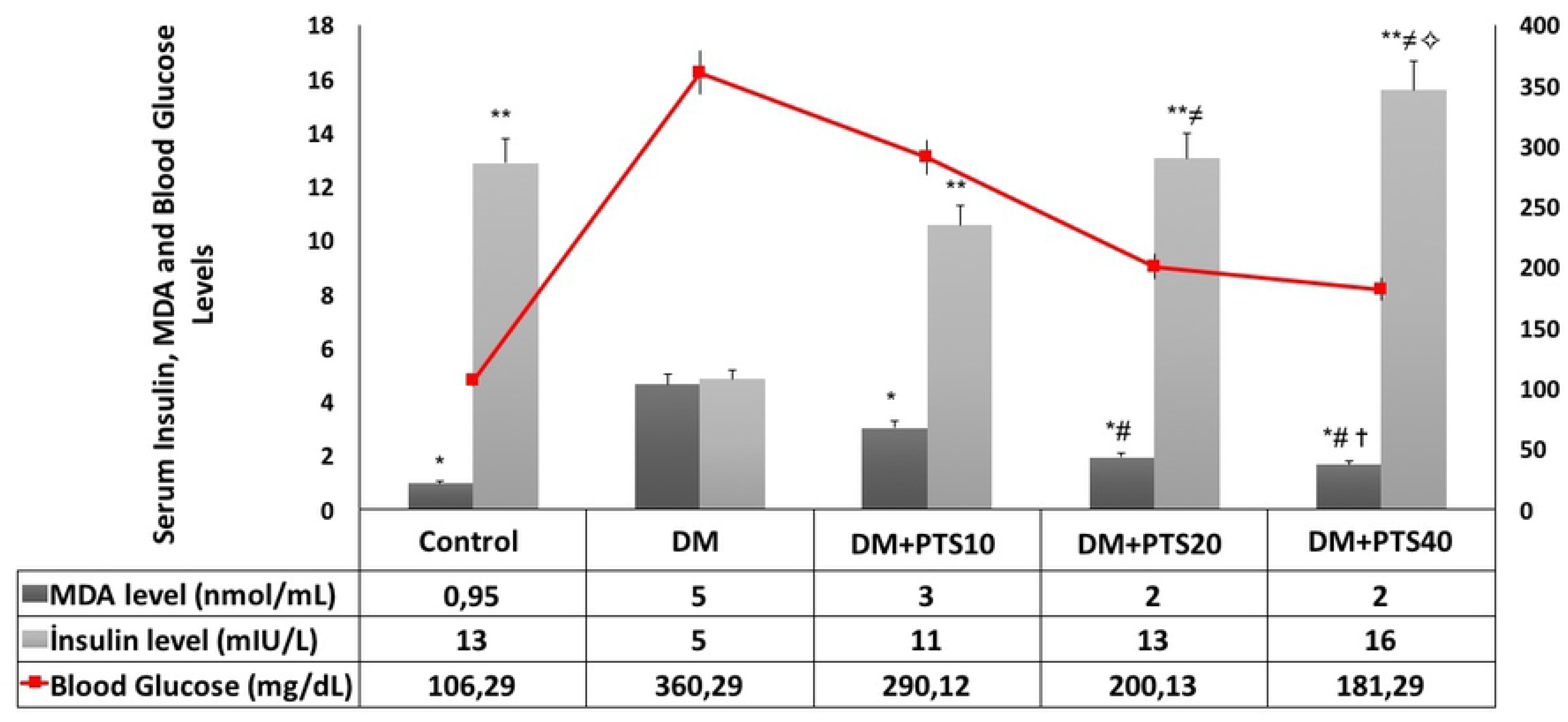
The serum MDA, insulin and blood glucose level change of the rat groups. For serum MDA values, ∗ significantly different from DM, # significantly different from DM+PTS10 (P< 0.001 for all), and ✝similar to Control (P= 0.075). For serum insulin values, ∗∗ significantly different from DM (P< 0.001 for all), ≠ different from DM+PTS10 (P< 0.001 for all), and ✧ similar to Control (P= 0.014). For blood glucose level; DM> PTS10 > PTS20 > PTS40 > Control (P< 0.001 for all).

The serum insulin levels were significantly lower in the untreated diabetic rats compared to the healthy Control group and all the PTS treatment groups (P < 0.001 for all). The diabetic rats that received PTS had increased insulin levels after 10, 20, and 40 mg/kg/day of PTS compared to the untreated diabetic rats (P < 0.001 for all). However, 20 and 40 mg/kg/day of PTS treatment were more effective in increasing serum insulin levels than the 10 mg/kg/day PTS dose (P < 0.001 for all) (Figure 7).

Serum MDA levels were significantly higher in the healthy Control group and in all the PTS treatment groups compared to the diabetic rats (P < 0.001 for all). The diabetic rats that received PTS (in all the treatment groups) had decreased serum MDA levels compared to the untreated diabetic rats (P < 0.001 for all). However, the PTS administrations of 20 and 40 mg/kg/day were more effective in lowering serum MDA levels in the diabetic rats than the lower treatment dose (P < 0.001 for all). The serum MDA levels of the healthy Control group rats were similar to those of the rats that received 40 mg/kg/day of PTS (P = 0.014) (Figure 7).

## Discussion

Our study aimed to investigate the effects of PTS, an antioxidant, on the cochleas of diabetic rats. Using an STZ-induced diabetic rat model, we found that PTS reduced serum MDA and blood sugar levels and increased insulin levels; in addition, PTS increased cell viability by inhibiting apoptosis in the cochleas. Our results showed that PTS protected the cochleas from DM-induced ototoxicity by increasing the DPOAE amplitude averages in a dose-dependent manner. Our study confirmed the anti-apoptotic effect of PTS in cochleas isolated from diabetic rats using the expression profiles of factors associated with apoptotic pathways. This data provides a basis for further studies.

DM is a metabolic disease with multiple etiologies. DM is characterized by metabolic disorders that are related to defects in insulin secretion and/or insulin resistance, and it is characterized by chronic hyperglycemia. DM is also associated with increased oxidative stress.^1^ DM induces oxidative and nitrosative stress, damaging DNA. Over time, DM becomes involved in the pathogeneses of cancer, the nervous system, and cardiovascular diseases.^3^ DM is a significant global public health problem, and there is a need for alternative prevention and treatment starategies.^1,2^

Recent research has shown that polyphenolic compounds have anti-diabetic potential and may prevent hyperglycemia.^27^ The polyphenolic-derivative PTS is a therapeutic agent with a high (80%) bioavailability, as its hydrophobic structure facilitates intracellular uptake.^28^ Research has shown that PTS’s antioxidant properties have protective and therapeutic effects in many diseases, including neurological, cardiovascular, diabetic, dyslipidemic, and hematological disorders.^17,29,30^ In addition, PTS has been reported to have anti-inflammatory effects,^31^ restorative immune system effects,^32^ and potent anti-cancer effects.^33,34^ Although studies have demonstrated the benefits of PTS for DM, the reports in the literature regarding the effect of PTS on the cochlea are so far inadequate.^29,35^

Sensorineural hearing loss is associated with several etiological factors, including congenital diseases, ototoxic agents, infectious diseases, and metabolic diseases, such as DM.^9^ Increased oxidative stress induced by DM can cause hearing disorders by accelerating neuron loss and demyelination in neuronal tissues, leading to peripheral endoterial dysfunction and microvascular complications.^4,5,10^ Studies have reported that DM causes various histopathological changes in the cochlea.^10^ Wackym, Linthicum and Smith et al. reported that spiral ganglion neurons in the cochlea deteriorated due to diabetic microangiopathy.^37,38^ In a human temporal bone study, Fukushima et al. reported that DM causes the loss of outer hair cells, a thickening in the basal membrane of the stria vascularis and the Corti organ, and atrophy in the stria vascularis.^39^ In a study with genetically diabetic rats, Nakae et al. observed stria vascularis degeneration, edema, and the loss of inner-outer hair cells on the basilar membrane.^40^ However, Tachibana-Nakae and Nageris et al. reported that there was no difference between the controls and the diabetic animals in terms of cochlear histopathology.^41,42^

The TUNEL assay provides an objective indicator of non-specific cell deterioration.^43^ Our histopathological analysis revealed significant TUNEL staining in the inner and outer hair cells of the Corti organs of the diabetic rats compared to the rats in the Control group. However, the rats in the PTS treatment group had decreased dose-dependent TUNEL-reactivity, likely because the PTS treatment protected the cochleas from DM-induced ototoxicity. In addition, apoptotic factors related to several apoptotic pathways were observed in immunohistochemical staining. There was significant caspase-8 and cytochrome-c immunostaining in the inner and outer hair cells of the Corti organs of the diabetic rats compared to those of the Control group.^44^ However, increased PTS doses decreased caspase-8 and cytochrome-c expression in the same cell types. The intrinsic apoptotic pathway is controlled by the mitochondrial permeability of the pro-apoptotic factor Bax and the anti-apoptotic factor Bcl-2 that is bound to caspase-3. Increased mitochondrial permeability and depolarization triggers cytochrome-c release, activating the apoptotic pathway related to caspase-3.^44,45^ In our study, PTS therapy in diabetic rats caused a dose-dependent decrease in the mRNA expression of pro-apoptotic *BAX* and its effector *CASP-3* and an increase in the mRNA expression of anti-apoptotic *BCL2*. The mRNA expression data of these pro- and anti-apoptotic genes correlate with the histopathological and immunohistochemical data, suggesting that an anti-apoptotic molecular mechanism is associated with the antioxidant PTS.

DPOAE is an important objective test used to determine hearing loss caused by damage to the outer hair cells in the cochlea. Cochlear deterioration may be caused by metabolic diseases, such as hyperlipoproteinemia, and DM has been reported to cause changes in DPOAE amplitudes.^25,46^ In our study, the mean DPOAE amplitudes were low in the diabetic rats and increased in a dose-dependent manner with PTS treatment. These data indicate that PTS may protect the outer hair cells, a finding that is consistent with our histopathological, immunohistochemical, and molecular findings, which all demonstrated decreased cochlear damage related to DM. Further studies with functional analyses are needed to assess the impact of PTS on the retrocochlea and the cochlea in the auditory pathway of diabetic animals.

The animal model for STZ-induced DM is characterized by elevated blood sugar levels, decreased serum insulin levels, and increased weight loss.^47^ Our study showed increased weight loss in untreated diabetic rats and decreased weight loss in the diabetic rats that received PTS treatment. Compared to the other doses, 40 mg/kg/day of PTS was associated with the least amount of weight loss. Our findings are in accordance with those of Korusu et al. and Sun et al.^48,49^ We believe that PTS reduces weight loss by promoting glycolysis, improving glycemic control, and maintaining adipose-muscle tissue. In our study, serum insulin levels were significantly lower in the diabetic rats compared to the healthy Control group and the treatment groups, likely due to the destruction of pancreatic beta-cells in the STZ-induced diabetic rats.^47,50^ In accordance with the literature, PTS treatment increased serum insulin levels in a dose-dependent manner. PTS may increase insulin sensitivity and increase hepatic hexokinase activity in glycolysis.^48,51^ In our study, PTS treatment caused a significant decrease in the blood glucose levels of the diabetic rats, particularly at doses of 20 and 40 mg/kg/day. These data are consistent with the work of Pari et al. and Manickam et al. and can be explained by the cytoprotective action of PTS, which promotes pancreatic beta-cell granulation.^19,35^ Previous studies have reported that MDA, a byproduct of lipid peroxidation, is an important oxidative stress marker, and that serum MDA levels increase in diabetic rats.^52,53^ Similarly, in this study, all PTS treatment doses decreased serum MDA levels, and this decrease was more pronounced at increased PTS doses. In our study, the decreased ototoxicity and increased cytoprotectivity of PTS in the cochleas of STZ-induced diabetic rats demonstrated the remedial effect of PTS on metabolic parameters.

## Conclusion

PTS can be used to control blood glucose and serum insulin levels and to reduce oxidative stress in STZ-induced diabetic rats. There was a significant increase in the dose dependent mean DPOAE values of the PTS-treated diabetic rats compared to the untreated diabetic rats. There was also a significant decrease in the number of apoptotic cells in the cochleas of the PTS-treated diabetic rats compared to those of the untreated diabetic rats. PTS had a significant dose-dependent, protective effect against DM-induced ototoxicity by inhibiting the intrinsic apoptotic pathway in the treatment group. The biochemical data in our study, showing that PTS has a dose dependent effect in STZ-induced diabetic rats, are consistent with our cochlear histopathological, immunohistochemical, molecular, and auditory findings. This study provides the first findings indicating that PTS can prevent DM-induced ototoxicity and suggests that PTS should be considered as a potential therapeutic agent.

## Acknowledgments

Thanks to Sabinsa Corporation for supplying the PTS. This study was financially supported by the Celal Bayar University Scientific Research Projects Coordination Unit.

## Author Contributions

Concept - S.Ö.; Design – S.Ö, T.Ö.; Supervision - T.Ö.; Resources - B.T; Materials – T.Ö., B.T. S.G.G; Data Collection and/or Processing – S.E, B.T.; Analysis and/or Interpretation – S.G.G, O.G.; Literature Search – S.Ö, T.Ö.; Writing Manuscript - S.Ö.B.T; Critical Review - A.P.

